# Poly(ADP-ribose)-binding and macroH2A mediate recruitment and functions of KDM5A at DNA lesions

**DOI:** 10.1101/2020.07.27.223131

**Authors:** Ramhari Kumbhar, Jullian Perren, Fade Gong, David Corujo, Frank Medina, Andreas Matouschek, Marcus Buschbeck, Kyle M. Miller

**Author notes:** Correspondence. (K.M.M.).

## Abstract

The histone demethylase KDM5A removes histone H3 lysine 4 methylation, which is involved in transcription and DNA damage responses (DDR). While DDR functions of KDM5A have been identified, how KDM5A recognizes DNA lesion sites within chromatin is unknown. Here, we identify two factors that act upstream of KDM5A to promote its association with DNA damage sites. We have identified a non-canonical poly(ADP-ribose), (PAR), binding region unique to KDM5A. Loss of the PAR-binding region or treatment with PAR polymerase (PARP) inhibitors (PARPi) blocks KDM5A-PAR interactions and DNA repair functions of KDM5A. The histone variant macroH2A1.2 is also specifically required for KDM5A recruitment and functions at DNA damage sites, including homology-directed repair of DNA double-strand breaks and repression of transcription at DNA breaks. Overall, this work reveals the importance of PAR-binding and macroH2A1.2 in KDM5A recognition of damage sites that drive transcriptional and repair activities at DNA breaks within chromatin that are essential for maintaining genome integrity.

**Summary:** The histone demethylase KDM5A demethylates H3K4 to promote repair and transcriptional responses at DNA breaks. We identified poly(ADP-ribose)-binding and macroH2A1.2 as modulators of KDM5A association with DNA damage sites, revealing how KDM5A engages DNA breaks within chromatin.

## Introduction

DNA is recurrently damaged by endogenous processes including transcription and replication, as well as by dysregulated proteins and chemical reactions within cells (Chatterjee and Walker, 2017; Lindahl and Barnes, 2000; Tubbs and Nussenzweig, 2017; Xia et al., 2019). Exogenous agents including ultraviolet light, chemicals and cancer therapies (i.e., radiation) also damage DNA (Ciccia and Elledge, 2010; Jackson and Bartek, 2009). An inability to constrain and repair DNA damage can result in mutations and genome instability. Loss of genome integrity can have dire consequences for cellular and organismal homeostasis, including the development of cancer (Jeggo et al., 2016; Negrini et al., 2010). Cells employ an extensive network of proteins, referred to as the DNA damage response (DDR), which counteract DNA damage by sensing its presence in the genome, activating the appropriate response and repairing the lesion (Ciccia and Elledge, 2010; Jackson and Bartek, 2009). Mutations within DDR related genes are frequent in cancer (Knijnenburg et al., 2018), which helps explain the prevalence of genome instability in cancer genomes. For example, DNA double-strand breaks (DSBs) are repaired mainly by homologous recombination (HR) and non-homologous end-joining (NHEJ) (Chapman et al., 2012), with the HR factors BRCA1 and BRCA2 found recurrently mutated in breast and ovarian cancer (Stratton and Rahman, 2008). While mutations in DNA repair genes result in genome instability, these genetic alterations also provide vulnerabilities that can be targeted therapeutically in some genetic backgrounds (Ashworth and Lord, 2018; Jackson and Helleday, 2016; O’Connor, 2015; Pilie et al., 2019). This strategy has been successfully employed against BRCA-deficient cancers, which display synthetic lethality with poly(ADP-ribose) polymerase (PARP) inhibitors, a treatment now used clinically (Bryant et al., 2005; Farmer et al., 2005; Fong et al., 2009; Lord and Ashworth, 2017).

Chromatin is an integral component of the DDR as DNA is organized by histones into chromatin, thus controlling the accessibility and their functions there (Agarwal and Miller, 2016; Caron et al., 2019; Chiu et al., 2017; Clouaire and Legube, 2019; Dabin et al., 2016; Gong and Miller, 2019; Kim et al., 2019c; Tan and Huen, 2020). One such chromatin mechanism involved in DDR regulation is histone post-translational modifications (PTMs). Histone marks including phosphorylation, methylation, acetylation and ubiquitylation act to regulate chromatin structure and function to coordinate the DDR with DNA-templated processes such as transcription, replication and repair (reviewed in; (Kim et al., 2019c). Histone variants, including the histone H2A family variants H2AX, H2AZ and macroH2A also participate in the regulation of the DDR (Buschbeck and Hake, 2017; Corujo and Buschbeck, 2018; Kim et al., 2018; Turinetto and Giachino, 2015). Collectively, chromatin employs several mechanisms to engage DDR factors to facilitate DNA damage signaling and repair within the chromatin environment.

Histone methylation plays important roles in the DDR (Gong and Miller, 2019; Kim et al., 2019c). As an example, lysine (K)-specific demethylase 5 (KDM5) subfamily proteins, which consist of four jmjC proteins, KDM5A-D, target the transcription-associated marks H3K4me2/3 to regulate transcription and the DDR, such as DSB repair, p53 regulation and the response to oxidative stress (Batie et al., 2019; Bayo et al., 2018; Christensen et al., 2007; Gong et al., 2017; Hu et al., 2018; Klose et al., 2007; Li et al., 2014b; Xu et al., 2018). KDM5A is recruited to DSBs where it demethylates H3K4me3 to regulate the ZMYND8-NuRD (nucleosome remodeling and histone deacetylation) chromatin remodeling complex (Gong et al., 2017). ZMYND8 binds acetylation marks at DSBs through its acetyl-binding bromodomain and recruits NuRD to repress transcription and promotes HR repair at DSBs (Gong et al., 2015; Gong et al., 2017; Gong and Miller, 2018; Savitsky et al., 2016; Spruijt et al., 2016). KDM5A is also found to be dysregulated in several cancers. For example, KDM5A is up-regulated in multiple cancers (Cao et al., 2014; Choi et al., 2018; Li et al., 2014a; Teng et al., 2013; Zeng et al., 2010) and promotes resistance to cancer treatments, including to DNA damaging agents (Hou et al., 2012; Sharma et al., 2010; Vinogradova et al., 2016; Yang et al., 2019). KDM5B is also involved in cancer and radioresistance (Bayo et al., 2018; Xhabija and Kidder, 2019). These findings have highlighted KDM5A, and other KDMs, as potential therapeutic targets in cancer. However, the potential connection between the DNA repair activities of KDM5A and its alterations in cancer have not been investigated. Although the function of histone demethylation by KDM5A in transcription and DNA damage repair have been described, the factors that regulate KDM5A in these processes are poorly understood.

Here, we have identified two mechanisms that regulate KDM5A localization to DNA damage sites. First, we identify a PAR interaction domain (PID) within KDM5A in its C-terminus that is required to recruit KDM5A to DNA damage sites. This domain is unique to this KDM5 lysine demethylase. KDM5A-PID by itself can localize to damage sites and its presence in KDM5A is required for its genome integrity promoting functions. Second, the histone H2A variant, macroH2A1.2 promotes KDM5A recruitment to DNA damage. Functionally, loss of macroH2A1.2 results in defective HR and DSB-induced transcriptional repression, which phenocopies KDM5A deficiency. Thus, we propose that PAR-binding and macroH2A1.2 recognition engage KDM5A to target this enzyme to DNA damage sites. Identification of KDM5A regulators in the DDR may provide valuable insights into their potential involvement in other biological processes dependent on this lysine demethylase, including transcription and cancer.

## Results and discussion

### KDM5A promotes genome integrity

We previously demonstrated the involvement of KDM5A in regulating the ZMYND8-NuRD chromatin remodeling complex at DNA damage sites. The KDM5A-ZMYND8-NuRD axis functions to demethylate H3K4me3 at DNA damage sites, to allow ZMYND8-NuRD to localize to DNA lesions within transcriptionally active chromatin, where this pathway represses transcription and promotes homologous recombination (HR) (Gong et al., 2015; Gong et al., 2017; Gong and Miller, 2018; Savitsky et al., 2016; Spruijt et al., 2016). To gain further insights into the involvement of KDM5A in genome integrity pathways, we obtained colon carcinoma cancer cell lines lacking KDM5A, which were generated from HCT116 cells using CRISPR-Cas9 gene editing. To control for any potential off-target gene editing effects, we reconstituted KDM5A knockout (KO) HCT116 cells with GFP-tagged full-length KDM5A (Fig. S1 A). Consistent with the involvement of KDM5A in HR, cells deficient for KDM5A were sensitive to the PARP inhibitor Olaparib (Fig. 1 A; KDM5A protein levels of cells in A analyzed in Fig. S1 A). PARPi sensitivity often correlates with HR deficiency as cells deficient for HR factors, such as BRCA1 or BRCA2 are sensitivity to PARP inhibition (Bryant et al., 2005; Farmer et al., 2005; Lord and Ashworth, 2017; O’Connor, 2015). The observed sensitivity of KDM5A KO cells to PARPi was largely rescued by complementation with ectopically supplied KDM5A, confirming that these results were due to loss of KDM5A in these cells (Fig. 1 A). PARP1 activity is required for KDM5A localization to the DNA damage sites to allow for efficient DSB repair by HR (Gong et al., 2017). Similar to KDM5A KO cells, human U2OS cells treated with the KDM5A inhibitor CPI455, which blocks the catalytic activity of this enzyme, displayed reduced cell survival after PARPi treatment compared to untreated cells or cells treated with either single inhibitor (Fig. 1 B). KDM5A-deficient cells treated with PARPi exhibited several phenotypes associated with the loss of genome integrity pathways, including increased micronuclei formation and persistence of DNA double-strand breaks as detected by the DNA damage marker histone variant H2AX phosphorylation (γH2AX) and neutral comet assays (Fig. 1 C-F; and Fig. S1 B-C). Loss of KDM5A did not alter the expression of other JARID1A histone demethylases, KDM5B and KDM5C, suggesting these results are specifically due to KDM5A deficiency (Fig. S1 D). These results provide strong evidence for the role of KDM5A in genome integrity pathways, including those that are affected by PARPi (i.e. HR repair).

**Figure 1.**
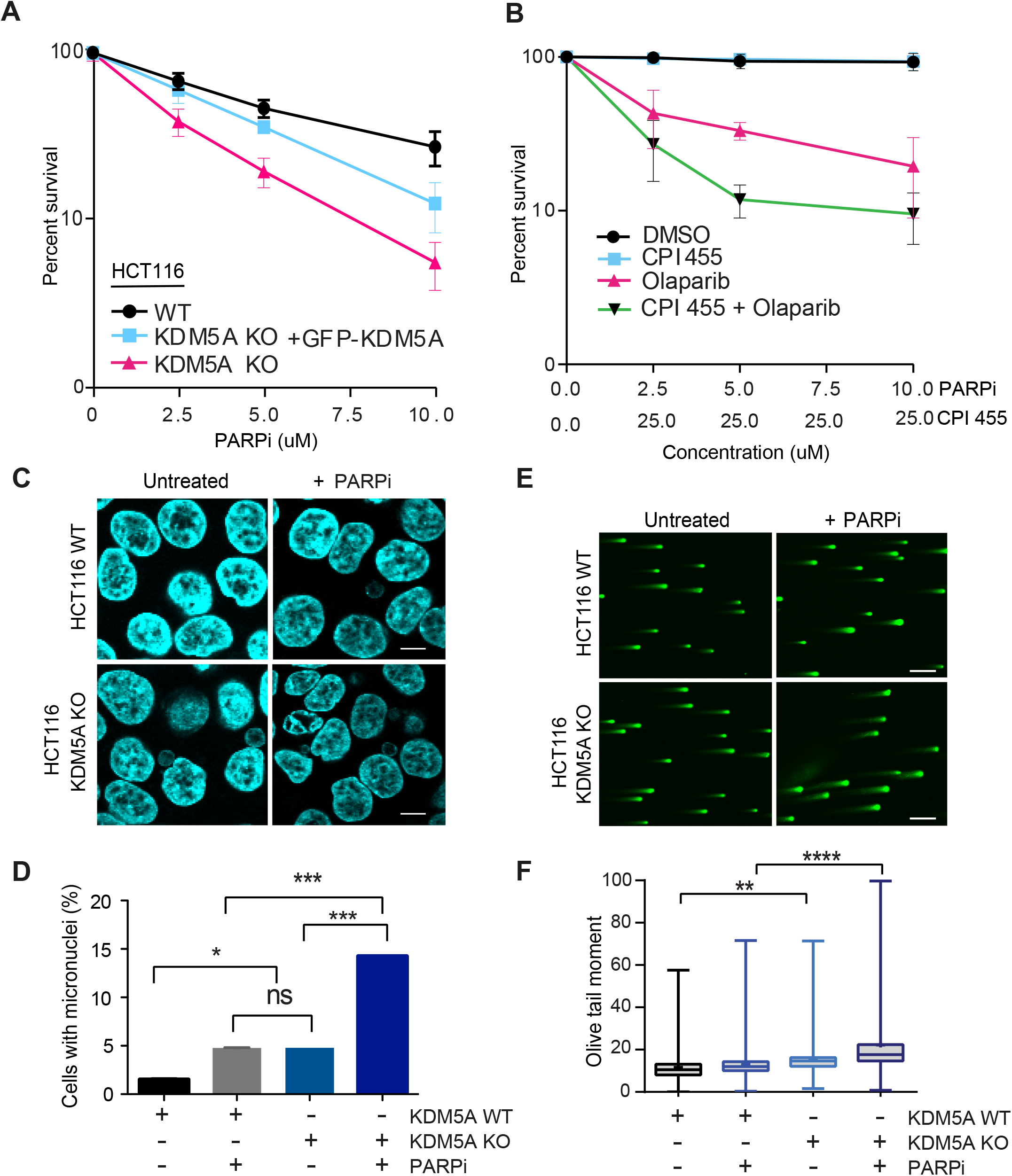
KDM5A deficiency results in genome instability and sensitivity to PARP inhibition. **(A)** Clonogenic survival assay of HCT116 WT, KDM5A KO and KDM5A KO + GFP-KDM5A complemented cells. Cells were treated with PARPi, Olaparib at the indicated doses for 24 h, and colonies were quantified after 2 weeks; Error bars represent SD; n=3. **(B)** Clonogenic survival assay of U2OS cells treated with DMSO, 25 μM CPI455 and Olaparib at indicated doses for 24 h. Colonies were analyzed as in A. Error bars represent SD; n=3. **(C)** KDM5A suppresses micronuclei formation. HCT116 WT and KDM5A KO cells were treated with 5 μM Olaparib for 24 h and immunostained with DAPI to detect micronuclei (ex. arrows). **(D)** Quantification of C. >100 cells quantified per condition; n=2. Scale bars, 10 μm. **(E)** Loss of KDM5A increases DSBs as detected by neutral comet assay. Cells were treated as in D. Scale bars = 100 μm **(F)** Quantification of E. Olive tail moment for >100 cells quantified per condition; n=2. Error bars represent SEM. P-values were calculated by Turkey’s multiple comparison test (*, P<0.05; **, P<0.01; ***, P<0.001; ****, P<0.0001; ns, not significant).

### KDM5A interacts with PARP1 and poly(ADP-ribose) chains in cells

PARP1 recruitment and formation of PAR chains at sites of DNA damage is one of the earliest responses to DNA damage. PARylation promotes the rapid accumulation of DNA damage factors to DNA lesions, including DSBs (Ray Chaudhuri and Nussenzweig, 2017). To further delineate how PARP1 regulates KDM5A in the DDR, we tested whether KDM5A interacts with PARP1 directly by performing co-immunoprecipitation (Co-IP) followed by western blot analysis. IP of endogenous KDM5A with an anti-KDM5A antibody in HEK293T cells indicated an interaction between endogenous PARP1 and KDM5A (Fig. 2 A). Reciprocal Co-IP western blot analysis using PARP1 antibodies in HEK293T cells corroborated the interaction between KDM5A and PARP1 (Fig. 2 B). The interaction between PARP1 and KDM5A was still detected and not increased in cells treated with ionizing radiation (IR) compared to cells that were undamaged (Fig. 2 A-B). Reduced PARP1-KDM5A interactions upon IR treatment could be due to increased binding of PARP1 to DNA or DNA damage induced autoPARylation of PARP, behaviors of PARP1 which are known to disrupt protein-protein interactions (Thomas and Tulin, 2013).

**Figure 2.**
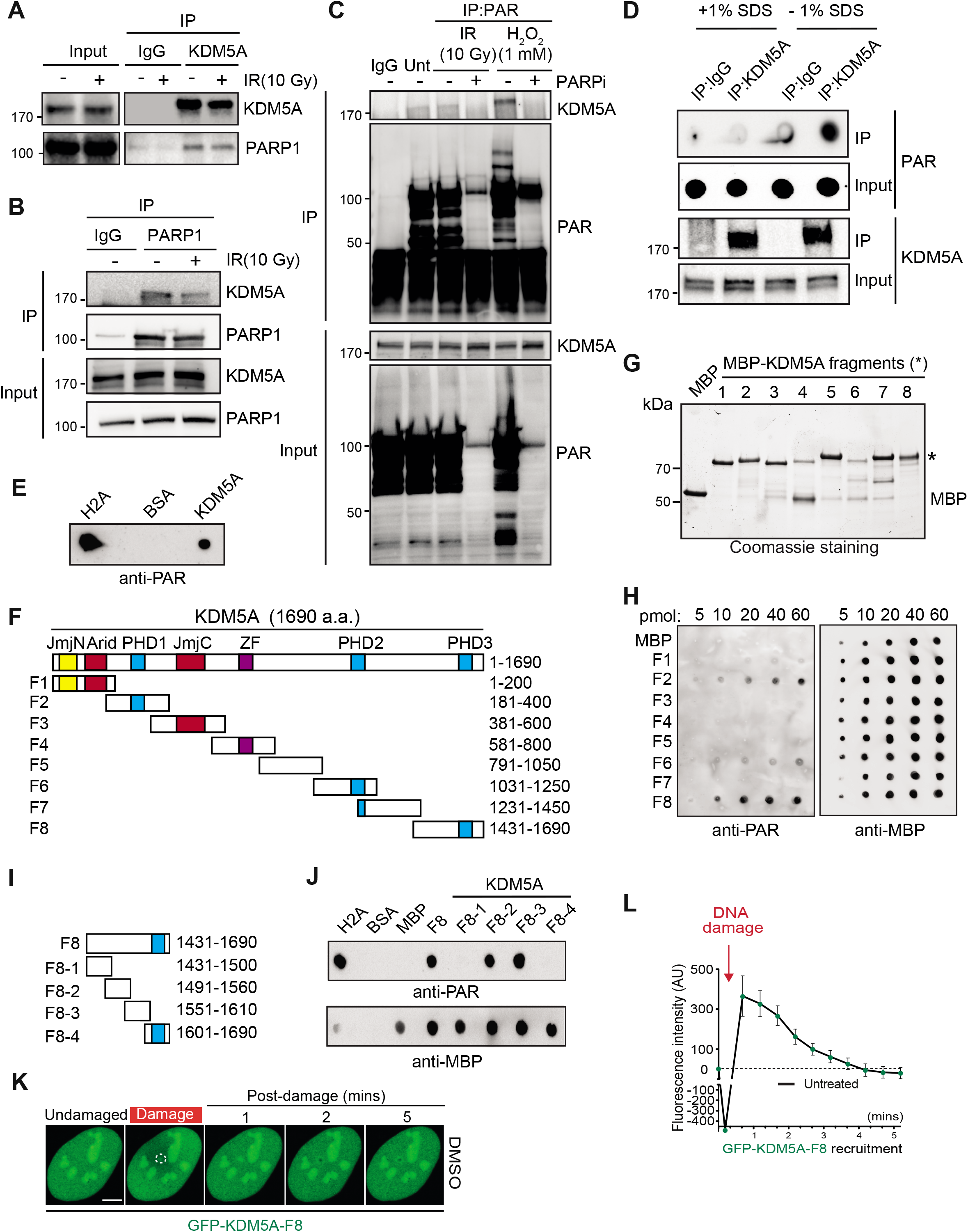
KDM5A interacts with PARP1 and Poly(ADP)-ribose chains. **(A)** Endogenous KDM5A and PARP1 interact. KDM5A was immunoprecipitated from HEK293T cells ± 10 Gy IR. Samples were analyzed by Western blotting with the indicated antibodies. **(B)** Reciprocal IP with endogenous PARP1 was performed as in A. **(C)** KDM5A binds to PAR in cells in a PARP-dependent manner. HEK293T cells were treated with Olaparib (5 μM, 1 h) followed by 10 Gy IR or H_2_O_2_ (1mM, 15 min) and examined by IP-Western with an anti-PAR antibody. **(D)** KDM5A binds PAR non-covalently. U2OS cells were treated with 10 Gy IR for 2 min, and cell lysates were incubated with KDM5A or IgG antibodies ± 1% SDS. Purified samples were either blotted on nitrocellulose membrane to detect PAR chains or resolved on SDS-PAGE and immunoblotted to detect KDM5A. **(E)** Purified KDM5A from U2OS cells binds PAR chains *in vitro*. Purified proteins were spotted onto nitrocellulose and probed with purified PAR chains with binding detected with an anti-PAR antibody. **(F)** Domain organization of KDM5A. Schematics for overlapping KDM5A fragments (F1 to F8) are indicated. **(G)** Expression of affinity purified MBP-tagged KDM5A fragments. Purified fragments were analyzed by SDS-PAGE and Coomassie staining. **(H)** Purified MBP and MBP-KDM5A fragments F1 to F8 were analyzed for PAR binding as in E. **(I)** Schematics for KDM5A F8 and derivatives. Blue box indicates PHD3 domain. **(J)** PAR binding resides in KDM5A-F8-2 and F8-3. PAR binding was performed as in E with indicated proteins. **(K)** GFP-KDM5A F8 sufficient for recruitment to laser damage. Laser micro-irradiation was performed in U2OS cells and damaged region is indicated by dotted white circle. **(L)** Quantification of K from one representative experiment. N ≥10 cells. Error bars represent SEM. AU, arbitrary units. Scale bars, 5 μm.

Given that PARP1 catalytic activity promotes the recruitment of KDM5A to laser-induced DNA damage sites (Gong et al., 2017) and poly ADP-ribosylation nucleates the localization of many proteins in the vicinity of DNA damage sites (Gibson and Kraus, 2012; Liu et al., 2017; Ray Chaudhuri and Nussenzweig, 2017), we next assessed if KDM5A was associated with PAR in cells. Immunoprecipitation of KDM5A or anti-PAR antibodies revealed an association of PARylated proteins with KDM5A (Fig. 2 C; and Fig. S1 E), suggesting that KDM5A or its associated proteins are PARylated and/or bind poly(ADP-ribose) (i.e PAR) chains. The increased binding between KDM5A and PAR was further enhanced by treatment with H_2_O_2_, another DNA damage-inducing agent and potent activator of PARP1 (Fig 2 C and Fig. S1 F). Treatment of cells with the PARPi Olaparib inhibited KDM5A-PAR interactions (Fig. 2 C), further supporting that these interactions are PARP-dependent. These results established a connection between KDM5A and PARylation in cells but do not distinguish between PAR binding or direct PARylation of KDM5A and/or an interacting protein. To address these various possibilities, KDM5A was immunoprecipitated under denaturing conditions using 1% SDS and the presence of PAR chains was evaluated by western blotting. KDM5A-PAR interactions were readily detected under native conditions (i.e. in the absence of SDS) and were abolished by the addition of SDS to denature the samples, which suggested KDM5A was not itself PARylated (Fig. 2 D). These interactions are also likely not mediated by nucleic acids given that cell lysates were treated with a broad spectrum DNA/RNA nuclease. Collectively, these results show that KDM5A interacts with PARP1 and PAR chains in cells in response to DNA damage.

### KDM5A directly binds PAR chains *in vitro*

Our results could be explained by the direct binding of KDM5A to PAR chains as a mechanism to direct its association to DNA damage lesions. To test this hypothesis, we determined if KDM5A was capable of binding PAR chains directly using *in vitro* PAR binding assays (Ahel et al., 2008; Kumbhar et al., 2018). Purified target proteins, including IP-ed KDM5A, histone H2A (a strong PAR binding protein) and BSA (negative control), were blotted onto nitrocellulose membrane and probed with purified PAR chains. Using purified KDM5A from U2OS cells, we observed KDM5A binding to PAR chains (Fig. 2 E and S1 G). These results were consistent with KDM5A binding directly to PAR chains and prompted us to identify the PAR binding region within KDM5A. Several PAR binding domains have been reported (Krietsch et al., 2013; Teloni and Altmeyer, 2016; Wei and Yu, 2016) but none of these are annotated within KDM5A in PFAM. Thus, to identify the PAR-binding region of KDM5A, we dissected full-length KDM5A into 8 partially overlapping MBP-tagged fragments (F1-F8), ensuring that the main functional domains of KDM5A were preserved within the fragments (Fig. 2 F). These fragments were expressed and purified from *E. coli*, then used in *in vitro* PAR binding assays (Fig. 2 G). These experiments identified 3 putative PAR binding regions, and the C-terminal fragment of KDM5A (F8), which contains amino acids 1431-1690, exhibited the strongest binding to PAR chains (Fig. 2 H and S1 H).

KDM5A-F8 fragment contains the third PHD domain of KDM5A, which interacts with its substrate H3K4me3 and is dispensable for supporting KDM5A localization to DNA damage sites (Gong et al., 2017; Wang et al., 2009). To further map the PAR interaction domain within KDM5A, we created four additional truncation mutations within fragment 8, including one containing the PHD3 domain (Fig 2 I). F8-2 (a.a. 1491-1560) and F8-3 (a.a. 1551-1610), displayed PAR binding and as expected, the fragment containing the PHD3 domain did not bind PAR (Fig. 2 J). These data show that the PAR binding domain resides within amino acids 1491-1610 in the C-terminus of KDM5A and importantly, is independent from the PHD3 domain.

We next sought to determine if the KDM5A-F8 region was sufficient to support DNA damage localization. To this end, we generated GFP-tagged KDM5A fragment 8 (GFP-KDM5A-F8) and analyzed its ability to be recruited to laser-induced DNA damage in cells. We observed rapid recruitment of GFP-KDM5A-F8 to DNA damage sites after micro-irradiation (Fig. 2 K, quantified in L). Retention of KDM5A-F8 was transient, with the maximum intensity of recruitment occurring within the first minute post-damage induction followed by a steady decline in localization at damage sites. These recruitment dynamics were reminiscent of the short lifespan of PAR chains, which are rapidly removed by Poly ADP-ribose hydrolase (PARG) (Alvarez-Gonzalez and Althaus, 1989; Kassab et al., 2020; O’Sullivan et al., 2019). In sum, these results map the PAR interaction domain of KDM5A within the amino acids 1491-1610 and show that this region is sufficient for localization to damage sites, supporting its identification as a PAR binding domain within KDM5A.

### Characterization of the PAR-binding domain in KDM5A

We next considered whether other KDM5 proteins interact directly with PAR chains. To address this question, we performed PAR-binding assay with purified GFP-KDM5A, GFP-KDM5B and GFP-KDM5C from U2OS cells. We did not include the male-specific KDM5D protein for this analysis. Although these proteins share similar domain structures and are highly homologous (Fig. 3 A), KDM5A, but not KDM5B and KDM5C, displayed PAR-binding activities (Fig. 3 B). While KDM5B has been reported to be involved in DSB repair in a PARP-dependent manner, our data may suggest that KDM5B is directly PARylated as a means to support its DNA damage functions (Li et al., 2014b). Taken together, these data suggest that direct binding to PAR chains by KDM5A is as unique mechanism that regulates its damage localization compared to other KDM5 demethylases.

**Figure 3.**
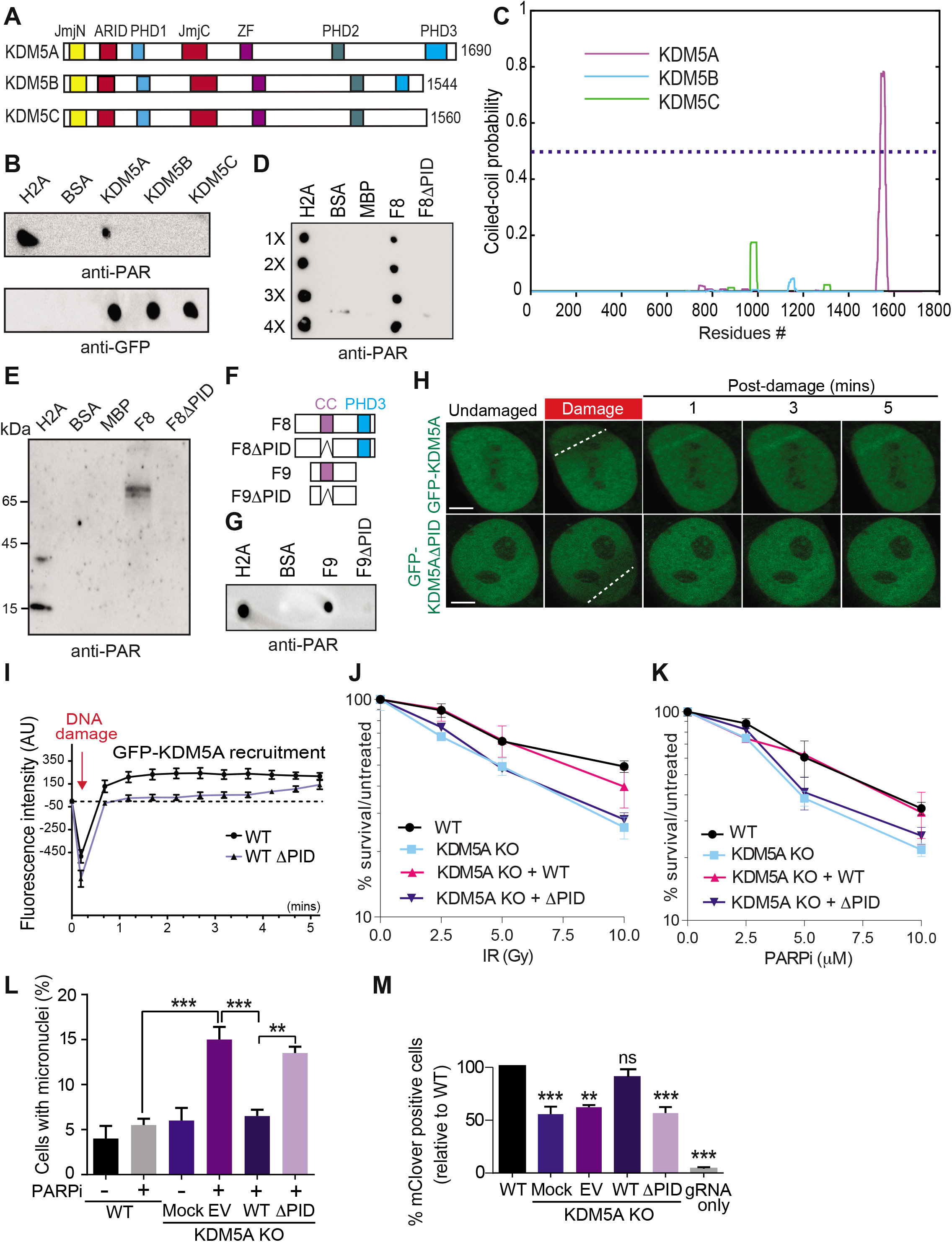
Identification and function of a poly(ADP)-ribose interaction domain within KDM5A. **(A)** Schematic of human KDM5A, KDM5B, and KDM5C. **(B)** KDM5A uniquely interacts with PAR chains. GFP-tagged KDM5A, KDM5B, and KDM5C were expressed and purified from U2OS cells. PAR binding was performed as in Fig. 2 E. **(C)** KDM5A contains a predicted coiled-coil region that is absent from KDM5B and KDM5C. Predicted per residue coiled-coil scores for dimers were calculated using Multicoil (Methods). **(D)** KDM5A region predicted as a coiled-coil (residues 1501-1562) is required for PAR binding (PID:PAR interaction domain). PAR binding assay performed as in B. **(E)** Far-Western analysis of KDM5A PAR binding. The indicated proteins were resolved by SDS-PAGE and blotted onto a nitrocellulose membrane. The membrane was incubated with PAR chains and the indicated proteins were detected as in Fig. 2 E. **(F)** Diagram of KDM5A-F8 and -F9 with putative coiled-coil (cc) and PHD3 domains indicated. **(G)** PAR binding analysis of KDM5A-F9 (1491-1610) and KDM5A-F9ΔPID. Protein purifications are described in material and methods. PAR binding assay was performed as in Fig. 2 E. **(H)** KDM5A PID is required for damage localization. Experiments were performed as in Fig. 2 K. Scale bars = 5μm **(I)** Quantification of H from one representative experiment. N≥10 cells. Damaged regions indicated by dotted white line. **(J-K)** KDM5A-PID is required for survival in response to IR and PARPi. Clonogenic survival assays were performed as in Fig. 1 A with the indicated ectopically expressed KDM5A genes; Error bars represent SD; n=3. **(L)** KDM5A-PID suppresses micronuclei formation. Experiments were performed as in Fig. 1E with indicated KDM5A KO cells and complemented cells as in J. (n=2), P-values were calculated by Turkey’s multiple comparison test (**, P<0.01; ***, P<0.001). **(M)** KDM5A-PID is required for efficient homology-directed repair (HDR). HDR efficiency in WT and KDM5A KO cells ± SFB-KDM5A and SFB-KDM5AΔPID was determined using CRISPR-mClover HR assay. GFP+ cells represent a repair event. %GFP+ cells were normalized to WT. Error bars represent SEM, n=2.

To further define the molecular properties of the PAR-binding domain in KDM5A, we performed a multicoil sequence analysis of KDM5A, KDM5B and KDM5C. Multicoil algorithm predicts coiled-coil location and oligomerization in protein sequences (Wolf et al., 1997). This analysis revealed that the PAR-binding region of KDM5A contains a highly probable coiled-coil domain, which is absent in KDM5B and KDM5C (Fig. 3 C). The region consists of highly basic and hydrophobic amino acids, and is predicted to be intrinsically disordered by IUPred, which is ideal for PAR binding (Fig. S2 A and B). This putative coiled-coil PAR-binding domain of KDM5A is not conserved between KDM5B and KDM5C but is highly conserved across KDM5A homologs from higher eukaryotes (Fig. S2 B and C). Intrigued by these observations, we further tested the ability of GFP-tagged KDM5B and KDM5C to be recruited to laser-induced DNA damage sites but could not detect damage association of these demethylases (Fig. S3 A-B). This result is consistent with another report that did not observe KDM5B recruitment to DNA damage after laser-microirradation (Li et al., 2014b). Since KDM5A and KDM5B are both reported to demethylate H3K4me3 at DNA damage sites, we also tested if KDM5B could be recruited to laser-induced DNA damage in cells lacking KDM5A. Even in cells lacking KDM5A, KDM5B was unable to associate with DNA damage sites to the extent detectable by this approach (Fig. S3 C), suggesting different mechanisms regulate these two demethylases in the DDR. To further demonstrate that the putative coiled-coil region containing the PAR interacting domain (residues 1501-1562) is required for PAR binding, we deleted this region from MBP-tagged KDM5A-F8 to create KDM5A-F8ΔPID. KDM5A and KDM5A-F8ΔPID were purified from *E. coli* and tested for PAR binding using an *in vitro PAR* binding assay. This analysis confirmed that the KDM5A-PID contained within fragment 8 was required for PAR binding as removal of the proposed PAR interacting domain abolished KDM5A-F8-PAR interactions (Fig. 3 D and E; purified proteins shown in Fig. S3 D). We had previously identified two overlapping fragments within fragment 8 of KDM5A that interacted with PAR (Fig. 2 J; fragments F8-2 and F8-3). We purified an untagged KDM5A 1491-1610 fragment (referred to as F9) containing these two domains along with a F9ΔPID mutant of KDM5A and performed PAR-binding assays (Schematic of KDM5A-F8 and -F9 constructs shown in Fig. 3 F). Results using these constructs were similar, showing PAR binding to be dependent on the PID of KDM5A (Fig. 3 G; purified proteins shown in Fig. S3 E). These analyses, along with our previous binding studies, support the presence of a PAR interaction domain (PID) within the C-terminus of KDM5A that is specific to this KDM5 demethylase family member and that this domain is functional given its high conservation among other eukaryotes.

### KDM5A-PAR binding is essential for KDM5A recruitment and function at DNA damage sites

To test for the functional importance of the PAR binding region in KDM5A that we identified, we analyzed the ability of KDM5A protein that lacks the PAR binding region to bind to sites of DNA damage and act in the DDR. Interestingly, GFP-KDM5A lacking the PAR-binding region (KDM5AΔPID) displayed reduced recruitment to DNA damage sites compared to full-length GFP-KDM5A (Fig. 3 H and I). Defects in homology-directed repair, increased micronuclei formation and reduced survival to IR and PARPi occur in KDM5A KO cells (Fig. 3 J and M). Expression of KDM5AΔPID was unable to rescue these deficiencies in KDM5A KO cells, unlike full-length GFP-KDM5A (Fig. 3 J and M). While we previously reported that siRNA-mediated depletion, or chemical inhibition of PARP1, reduced KDM5A localization to DNA damage sites and that KDM5A was required for HR repair (Gong et al., 2017), these results now reveal the molecular mechanisms that explain these observations. Taken together, these data demonstrate that KDM5A utilizes a non-canonical, putative coiled-coil, PAR binding domain in its C-terminus to bind PAR chains at DNA damage sites, which is required to orchestrate its functions in promoting genome integrity and HR repair.

### KDM5A interacts with the histone variant macroH2A1

Given the importance of KDM5A in chromatin-based DNA repair activities, we sought to test if other histone pathways may regulate KDM5A activities. We focused on the histone variant macroH2A as this variant can bind and be regulated by PARP and is involved in HR (Khurana et al., 2014; Ruiz et al., 2019; Timinszky et al., 2009). macroH2A1 has two splice variants, macroH2A1.1 and macroH2A1.2, which differ in 32 amino acids within the C-terminal macrodomain leading to the presence of a PAR binding domain in macroH2A1.1 but not in macroH2A1.2 (Kozlowski et al., 2018; Timinszky et al., 2009). macroH2A2 is expressed from a second, independent gene and does not bind to PAR but is ~80% identical with macroH2A1 (Posavec et al., 2013). Immunoprecipitation of SFB-tagged KDM5A from HEK 293T cells revealed an interaction between KDM5A and macroH2A1, including after DNA damage induction by IR (Fig. 4 A). Reciprocal IP analysis using endogenous macroH2A1 antibodies corroborated these results (Fig. S3 F). To determine which macroH2A1 variant KDM5A interacted with, we performed endogenous KDM5A IP in HEK 293T cells and immunoblotted with antibodies specific to macroH2A1.1 and macroH2A1.2. We observed that KDM5A interacted specifically with macroH2A1.2 (Fig. S3 G). To further validate these results, we performed Co-IP experiments with KDM5A and macroH2A using a HepG2 cell line where all macroH2A variants are stably knocked-down and complemented with stably expressed Flag-macroH2A1.2 or Flag-macroH2A2 (Douet et al., 2017). Co-IP western blot analysis using HepG2 control and macroH2A depleted derivatives identified an interaction with macroH2A1.2 but not macroH2A2 (Fig. 4 B). We next considered that the interaction between KDM5A-PARP1-macroH2A1.2 may be regulated by DNA damage. To this end, we performed Co-IP experiments in cells untreated or treated with IR in the presence or absence of PARPi. As we previously demonstrated, KDM5A interacted with PARP1 and macroH2A1.2 in undamaged or damaged conditions (Fig. 4 A and C). However, specifically in IR-treated conditions, these interactions were dependent on PARP activity as the addition of PARPi reduced the interaction between KDM5A and both PARP1 and macroH2A (Fig. 4 C). This is interesting given that macroH2A1.2 does not interact with PAR chains directly. We reasoned that PAR binding by KDM5A may mediate these interactions. To test this hypothesis, we asked if KDM5A-PID is required for KDM5A interactions with PARP1 and macroH2A1.2. Indeed, KDM5AΔPID, which lacks PAR binding at DNA damage sites, failed to associate with either macroH2A1.2 and PARP1 as efficiently as WT KDM5A (Fig. 4 D). These results identify KDM5A PAR-binding as being important for both PARP1 and mH2A1.2 interactions (summarized in Fig. 4 E).

**Figure 4.**
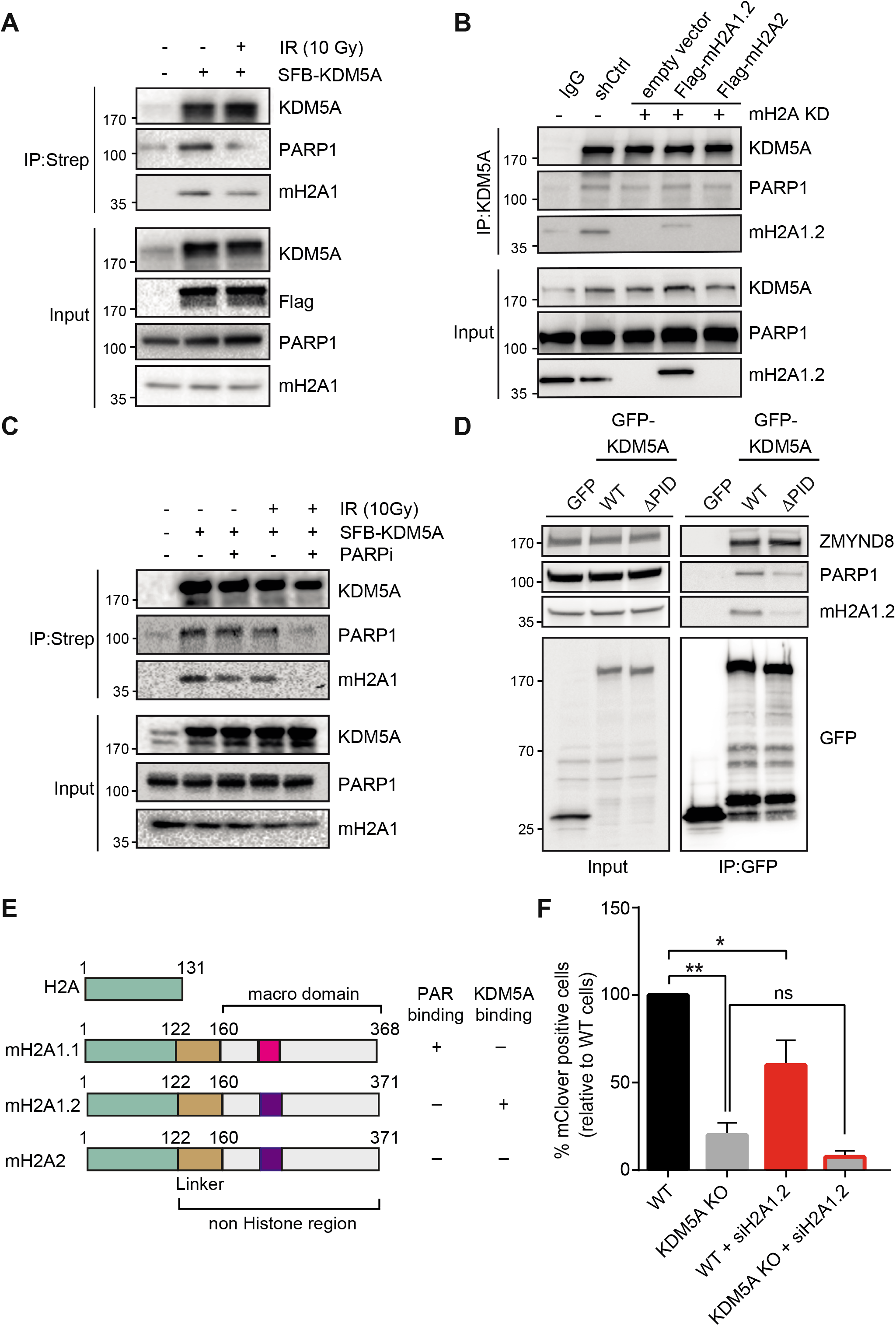
KDM5A interacts with the histone variant macroH2A1.2. **(A)** KDM5A interacts with macroH2A1. Inducible SFB-KDM5A expressing HEK293T cells were treated with ± 10 Gy IR. SFB-KDM5A was immunoprecipitated with streptavidin beads and interactions were detected by Western blotting. Input shows expression and loading of proteins. **(B)** Endogenous KDM5A interacts with macroH2A1.2. Co-IPs were performed using HepG2 cells depleted with shRNA control (shCtrl) or shRNA macroH2A (mH2A KD), and complemented shRNA-macroH2A with Flag-macroH2A1.2 or Flag-macroH2A2. **(C)** KDM5A and macroH2A1.2 interactions are PARP-dependent. SFB-KDM5A expressing HEK293T cells were treated with DMSO or Olaparib (5 μM, 1 h) followed by ± 10 Gy IR treatments. Co-IPs were performed as in A. **(D)** The PID of KDM5A is required for macroH2A1.2 and PARP1 interactions. HEK293T cells expressing GFP, GFP-KDM5A, or GFP-KDM5AΔPID were analyzed by Co-IP Western blot analysis with the indicated antibodies. **(E)** Comparison of domain structure of histone H2A and macroH2A variants with PAR and KDM5A binding summary. **(F)** KDM5A and macroH2A1.2 promote HDR. HR efficiency in WT and KDM5A KO cells ± siRNA depletion of macroH2A1.2 was performed as in Fig. 3 M; (n=2). Error bars represent SEM. P-values were calculated using an unpaired Student’s t-test (*, P<0.05; **, P<0.01; ns, not significant).

### Histone variant macroH2A1.2 is required for KDM5A recruitment to DNA damage sites

Histone variant macroH2A1.2 but not macroH2A1.1 has been reported to be required for efficient DSB repair through HR by promoting BRCA1 retention at DSBs (Khurana et al., 2014). Since macroH2A1.2 and KDM5A are associated with DSB repair by HR, and we identified an interaction between these proteins, we set out to determine if macroH2A1.2 and KDM5A collaborate to promote DNA repair. To begin to address this question, we measured HR efficiency using a CRISPR-Cas9/mClover assay in cells single or doubly deficient for KDM5A and macroH2A1.2 (Kim et al., 2019b; Pinder et al., 2015). In agreement with our previous results and other studies, we observed that KDM5A or macroH2A1.2 resulted in decreased gene-targeting efficiency, which is a readout for HR, compared to WT control cells (Fig. 4 F). Depletion of H2A1.2 in KDM5A KO cells did not further decrease HR efficiency significantly (Fig. 4 F).

Considering these results, we speculated that macroH2A may regulate the recruitment of KDM5A to DNA damage sites. We previously generated CRISPR-Cas9 gene-edited U2OS cells that lack macroH2A (Leung et al., 2018). Using these cells, we sought to determine if macroH2A is required for KDM5A recruitment to laser-induced DNA damage sites. To this end, we observed that GFP-KDM5A is recruited less efficiently to DNA damage sites in cells lacking macroH2A compared to WT cells (Fig. 5 A and B). We extended these results by analyzing the recruitment of ZMYND8, a protein that relies on KDM5A to support its accumulation at DNA damage sites through a H3K4me3 demethylation step in chromatin associated with DNA breaks and transcriptionally active loci (Gong et al., 2017). Indeed, macroH2A was also required for ZMYND8 damage recruitment (Fig. S3 H and I). Furthermore, we obtained similar results using HepG2 macroH2A-deficient cells for KDM5A. Importantly, reconstitution of these cells with macroH2A1.2 specifically led to a rescue of KDM5A accrual at damage sites (Fig. 5 C and D). Finally, we reasoned that if macroH2A1.2 was required in the KDM5A-ZMYND8 DDR axis, this histone variant would also be required to suppress transcription after DSB induction, a pathway reliant on KDM5A and ZMYND8. To test this idea, we used a well-established cell-based system to analyze transcriptional repression following DSB formation. This system contains a LacO array which is bound by an inducible LacI-FokI nuclease that creates DSBs upstream of a transcriptionally active gene (Fig. 5 E). Cells containing this system also express YFP-tagged MS2 which binds to the hairpin structures that occur within the mRNA that is driven by the transgene downstream of the LacO array. Upon DSB induction, the downstream gene is suppressed which halts production of the mRNA and abolishes the formation of colocalized mCherry-FokI and YFP-MS2 (Fig. 5 F and G). As expected, depletion of KDM5A resulted in defective transcriptional repression, causing an increased frequency of mCherry-FokI and YFP-MS2 foci. Depletion of macroH2A1.2 resulted in a loss of transcriptional repression, which phenocopied the loss of KDM5A (Fig. 5 F and G). These results demonstrate that macroH2A1.2 is required for break-induced transcriptional repression and suggest that macroH2A controls the KDM5A-ZMYND8 DDR pathway by promoting the recruitment of these factors to DNA damage sites.

**Figure 5.**
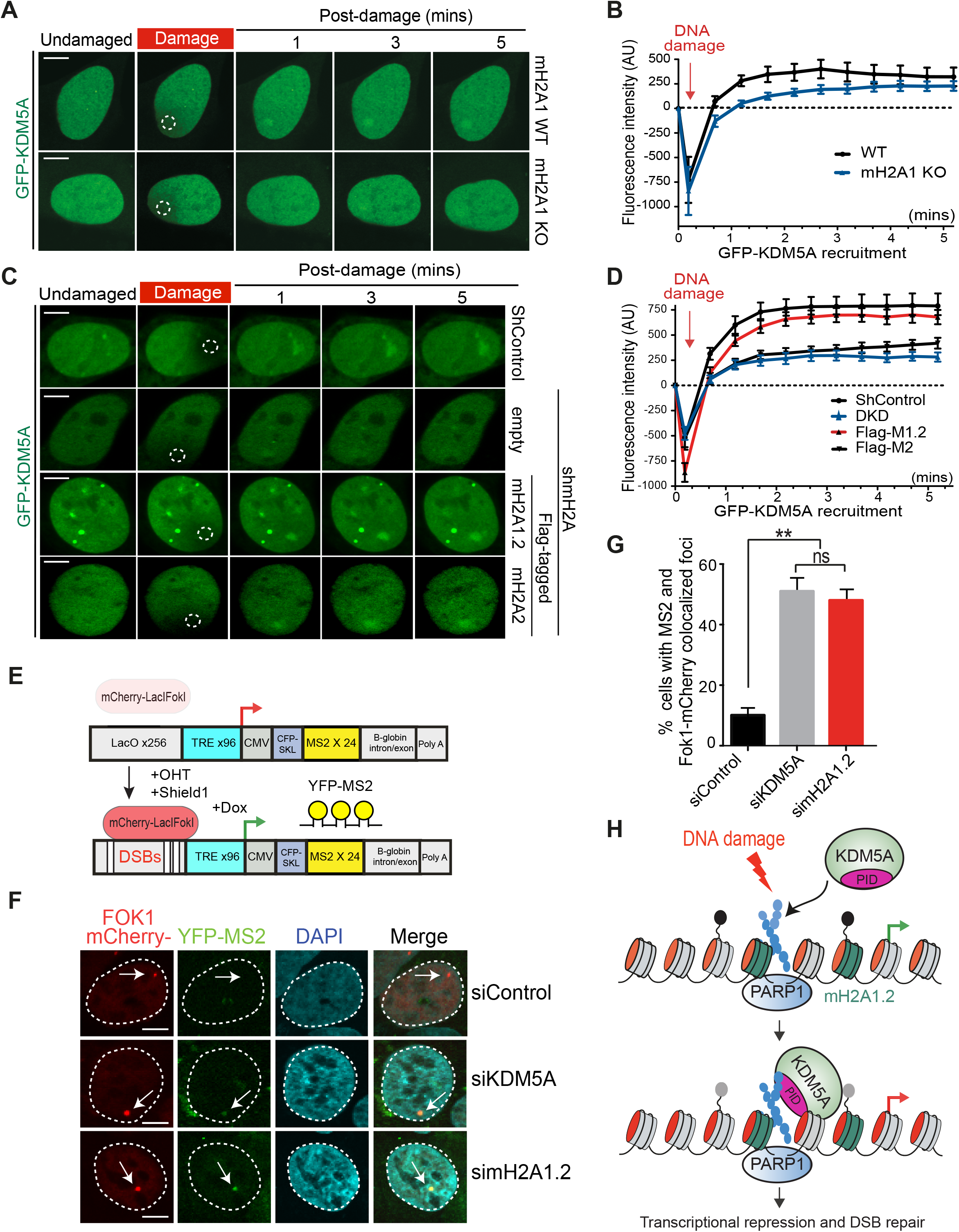
Histone variant macroH2A1.2 promotes KDM5A recruitment and transcriptional function at DNA damage sites. **(A)** macroH2A1 promotes KDM5A damage localization. Analysis of GFP-KDM5A damage recruitment in macroH2A1 KO U2OS cells was performed as in Fig. 3 H. **(B)** Quantification of A. n>10 cells. Error bars represent SEM. AU, arbitrary unit. **(C)** macroH2A1.2 promotes KDM5A damage recruitment. GFP-KDM5A damage recruitment was performed cells from Fig. 4 C. Dotted white circles indicate damaged regions. Scale bars = 5 μm. **(D)** Quantification of C as in B. n>10. **(E)** Schematic of FokI inducible DSB reporter cell system (Tang et al. 2013). Upon 4-OHT and Shield1 treatment, mCherry-FokI endonuclease induces DSBs upstream of a doxycycline inducible reporter gene. Nascent transcription is visualized by YFP-MS2 protein binding to stem loop structures in the mRNA. DSBs are visualized by mCherry-FokI localization to the LacO loci. **(F)** macroH2A1.2 is required for transcriptional repression following DSBs. Nascent transcription was analyzed at FokI-induced DSBs in siControl, siKDM5A and simacroH2A1.2 treated cells. Presence of MS2 foci indicates loss of DSB-induced transcriptional repression. **(G)** Quantification of F. >100 cells were analyzed per condition; Error bars represent SEM; n=2. **(H)** Model for KDM5A regulation by PARP1 and macroH2A1.2. Following DNA damage, PARP1 promotes PARylation and recruitment of KDM5A to DNA damage sites. KDM5A engages PAR chains through its C-terminal PID. macroH2A1.2 also promotes KDM5A accrual at DSBs to facilitate DSB-induced transcriptional repression and HR repair.

## Summary

Here, we report the identification of two molecular mechanisms that govern the DNA damage recruitment and function of KDM5A in the DNA damage response through the employment of cellular, genetic, and biochemical approaches. Our results reveal that KDM5A directly binds to PAR chains at DNA damage sites through a newly identified PAR interacting domain that resides within the C-terminus of KDM5A that is not found in other KDM5 demethylases. We also find that the histone variant macroH2A1.2 specifically interacts with KDM5A and promotes its recruitment to DNA damage sites. Our work thus identifies a PARP1-macroH2A1.2 pathway that acts upstream of KDM5A to control its interaction and functions at DNA break sites (Fig. 5 H).

Chromatin structure is known to be dynamically regulated upon DNA damage. Interestingly, PARP1 and macroH2A1.2 have opposing effects on chromatin structure. PARP1 and PARylation decompacts chromatin while macroH2A1 compacts chromatin, including through a mechanism involving PAR binding by the macrodomain of macroH2A1.1 (Khurana et al., 2014; Kozlowski et al., 2018; Krishnakumar and Kraus, 2010; Timinszky et al., 2009). macroH2A1.2 lacks a PAR-binding macrodomain but recent evidence has shown that this variant is still able to compact chromatin through an activity located in the linker region (Kozlowski et al., 2018). This bi-phasic behavior of chromatin decompaction followed by compaction is observed at DNA damage sites and has been shown to correlate with HR through a macroH2A1.2-dependent mechanism (Khurana et al., 2014). Our data demonstrating PARP1-macroH2A1.2 regulation of KDM5A may shed light on how these previous observations relate to KDM5A.

KDM5A is rapidly recruited to damage sites in a PAR-binding-dependent manner (Fig. 3 H). While the PAR binding region of KDM5A alone was rapidly recruited to damage sites, its association with breaks was very transient (Fig. 2 K and L). Transient PAR chains at breaks may be involved in the initial recruitment of KDM5A but additional mechanisms may be needed for retainment of these factors to DNA lesions. We posit that additional, multi-valent interactions may act to channel PAR-binding proteins into a specific pathway versus others that are regulated also by PARylation, for example singlestrand break repair, Okazaki fragment sensing and transcription (Gibson and Kraus, 2012; Hanzlikova et al., 2018; Ray Chaudhuri and Nussenzweig, 2017). Our data would suggest that macroH2A1.2 likely serves this purpose for KDM5A at DSBs, given that they interact with each other and the requirement of this histone variant for KDM5A recruitment and function at DNA damage sites.

Chromatin compaction at break sites may be in part related also to our observation that macroH2A1.2 is required for transcriptional repression at DNA breaks. Indeed, KDM5A demethylates the transcription active mark H3K4me3 at DSBs and promotes the damage association of ZMYND8-NuRD complex, which together act to repress transcription and promote HR repair of DSBs (Gong et al., 2015; Gong et al., 2017; Gong and Miller, 2018; Savitsky et al., 2016; Spruijt et al., 2016). Several other transcriptional repressive complexes, including PRC1, PRC2 and NELF (negative elongation factor), as well as the histone variant H2AZ, are also required for DNA break-induced transcriptional repression (Caron et al., 2019; Puget et al., 2019; Tan and Huen, 2020). Given the identification of the histone variant macroH2A1.2 in this damage response here, future studies are warranted to decipher other potential interactions between these DSB-associated transcriptional regulators.

PARP1 was one of the first factors identified that is required for break-induced transcriptional repression (Chou et al., 2010) and it was previously shown to regulate DNA damage recruitment of KDM5A (Gong et al., 2017). The identification of a PAR interaction domain in KDM5A was unexpected for several reasons. KDM5B, a related KDM5 demethylase, is also involved in DSB repair but is PARylated directly, which inhibits its activity at transcriptionally active sites (Bayo et al., 2018; Krishnakumar and Kraus, 2010; Li et al., 2014b). In addition, KDM5A does not share homology with any known PAR binding domains (Krietsch et al., 2013; Teloni and Altmeyer, 2016; Wei and Yu, 2016). Our extensive mapping and functional analysis identified a putative coiled-coil domain near the C-terminus of KDM5A able to bind PAR both *in vitro* and in cells as well as being sufficient to localize to DNA damage sites in a PARP-dependent manner. This strong evidence reveals that this domain can bind PAR and is necessary for KDM5A recruitment and activities at DNA damage sites. These findings could have several important implications. Coiled-coil domains are common in proteins, including several involved in the DDR, and play important roles in protein-protein interactions and protein multimerization and conformational changes (Truebestein and Leonard, 2016). In the DDR, dimerization of CtIP, a BRCA1 interacting protein requires a N-terminal coiled-coil region (Dubin et al., 2004), while BRCA1-PALB2 interaction is mediated by coiled-coil regions present in both proteins (Song et al., 2018). It will be important to test additional coiled-coil domains in other proteins for PAR binding, which may reveal additional mechanisms by which PARP1 regulates not only the DDR but also other PARP-dependent processes like transcription. Finally, KDM5A-PID is unique to this KDM5 demethylase. Given that KDM5A is overexpressed in cancer and is involved in therapy-associated drug resistance (Cao et al., 2014; Choi et al., 2018; Hou et al., 2012; Li et al., 2014a; Sharma et al., 2010; Teng et al., 2013; Vinogradova et al., 2016; Yang et al., 2019; Zeng et al., 2010), our data provide the rationale for testing the potential use of PARPi in these settings, which may selectively inhibit KDM5A.

In summary, this work has identified two regulatory mechanisms that control KDM5A interactions and function at DNA damage sites (Fig. 5 H). These findings exemplify how chromatin acts as a platform for various DNA templated processes, which must be remodeled by chromatin modifying enzymes to transition from one activity to another. In this example, upon DNA damage, PARP1 and macroH2A1.2 regulate chromatin structure and modifications to trigger KDM5A damage-association, resulting in histone demethylation and recruitment of the ZMND8-NuRD complex that collectively act to repress transcription and promote DNA repair.

## Materials and methods

### Cell lines and cell culture

Human osteosarcoma (U2OS) and human embryonic kidney (HEK293T) cells were purchased from ATCC, HCT116 WT and HCT116 KDM5A knockout cells were purchased from Horizon discovery. All cell lines were maintained in Dulbecco’s modified Eagle’s medium (DMEM) supplemented with 10% fetal bovine serum (FBS), 2 mM L-glutamine, 100 U/ml penicillin, and 100 μg/ml streptomycin at 37 C and 5% CO_2_. Inducible cell lines (SFB-tagged KDM5A), were established and maintained in medium with 0.2 mg/ml hygromycin B (Invitrogen). To establish the inducible SFB-tagged KDM5A expressing cell lines, pcDNA5/FRT/TO containing SFB-tagged constructs were transfected with pOG44 Flp Recombinase expression vector into Flp/In T-rex HEK293T cells. After 48 h, cells were treated with 0.2 mg/ml hygromycin B (Invitrogen) for the selection of transfected cells. All cell lines were routinely checked for mycoplasma contamination. Cells were pretreated with 5 μM PARP inhibitor (Olaparib), 5 μM PARG inhibitor PDD00017273 (Tocris) for 1 h before laser-induced DNA damage.

The U2OS DSB reporter cell line used for transcriptional analysis was a gift from Roger Greenberg, University of Pennsylvania (Tang et al., 2013). Cells were treated with 1 μM Shield1 and 1 mM 4-OHT for 3 h to induce site-specific DSBs and 1 μg/ml Dox for additional 3 h to induce transcription

The generation of HepG2 cells with stable shRNA-mediated knockdown of both macroH2A1 and macroH2A2 proteins is described in Douet et al., 2017. Cell lines stably expressing Flag-tagged macroH2A1.2 and macroH2A2 proteins in the double-knockdown background were achieved by retroviral transduction. GP2-293 cells were used as packaging cells to produce retroviral particles. Four million GP2-293 cells were seeded in P10 plates and cultured to 60-70% confluency, then transfected with 10 μg of pBabe.puro plasmids containing either Flag-tagged full-length mouse macroH2A1.2 or macroH2A2 sequences and 3 μg of pCMV-VSV-G mixed in a 1x HBS solution (2x HBS: 272 mM NaCl, 2.8 mM Na_2_HPO_4_, 55mM HEPES, pH 7) containing 125mM CaCl_2_. The supernatant containing viral particles produced by GP2-293 cells was collected at 24h and 48h after transfection, filtered using a 0.45 μm filter, supplemented with 8 μg/mL of Polybrene (Sigma-Aldrich) and added to the target HepG2 cells cultured in six-well plates at 60-70% confluency. Cells were then centrifuged 45 min at 1200rpm at 37°C, incubated at 37°C for 45min and then cultured o/n in fresh media. The same process was repeated 24h after the first infection. The cells were selected with 2 μg/ml puromycin. The necessary selection time was determined by using a negative control plasmid without resistance.

### Cloning and plasmids

KDM5A was cloned into the pDONR201 Gateway vector. The pDONR201 clones were transferred into GFP or SFB tagged DEST vectors using the Gateway LR cloning system (Invitrogen). KDM5AΔPID deletion mutant was generated by PCR in pDONR201 vector following standard cloning method and then subcloned into Gateway destination vector derived from pcDNA5/FRT/TO plasmids containing the NLS and SFB- or GFP-tagged in N-terminal. KDM5A fragments were amplified from KDM5A cDNA using Q5 High fidelity polymerase with primers containing *SalI* and *NotI* recognition sites. PCR products were then cloned into *SalI* and *NotI* sites of a pMAL-C5X vector (New England Biolabs). All constructs were validated by DNA sequencing.

### siRNA depletion and Overexpression

For knockdown experiments, 2.5×10^5^ cells were seeded in 6-well plates. Cells were transfected with siRNA duplexes targeting the corresponding protein or scrambled mixes using Lipofectamine RNAiMax (Invitrogen) reagents as per manufactures recommendations. Knockdown cells were cultured for 48-72 h before further processing. siRNA used are: KDM5A-5’-GCAAAUGAGACAACGGAAA-3’, macroH2A1.2 - 5’-CUGAACCUUAUUCACAGUGAA-3’,

For KDM5A rescue experiments or overexpression, cells were transfected with 2 μg of emGFP-KDM5A or emGFP-pcDNA 6.2 empty vector using either Lipofectamine 2000 or Lipofectamine 3000 and then allowed to recover for 24 h before proceeding to assay further. In the case of Dox inducible SFB- or GFP-tagged pcDNA5/FRT/TO plasmids containing KDM5A or KDM5AΔPID, 1 μg/ml Doxycycline was used for at least 24 h to induce expression of the indicated gene.

### Immunoprecipitation

Cells (U2OS or HEK293T) were lysed in NETN buffer (10 mM Tris-HCl, pH 8.0, 150 mM NaCl, 0.5% NP-40, protease inhibitor cocktail) containing TurboNuclease (Accelagen) at 4 °C for 1 h. Cell lysates were centrifuged at 15,000 rpm at 4 °C for 10 min. The lysate was then incubated with appropriate antibodies for 12 h and then conjugated with dynabeads Protein A or Protein G beads for an additional 1 h SFB- or GFP-tagged proteins were immunoprecipitated with streptavidin dynabeads (Invitrogen) or GFP-Trap (Chromotek). After 3-4 washes with NETN buffer, bead-bound proteins were eluted with 2X sample loading buffer and resolved on 4-15% gradient SDS-PAGE gel. Proteins were identified by an appropriate antibody by western blotting.

### Antibodies

Antibodies used in this studies are: mouse anti-KDM5A (ab78322; Abcam), rabbit anti-KDM5B (A301-813; Bethyl Laboratories, Inc.), rabbit anti-KDM5C (A301-034; Bethyl Laboratories, Inc.), rabbit anti-βtubulin (ab6046; Abcam), mouse anti-γH2AX (05-636 (JBW301); Millipore), rabbit anti-GFP (A11122; Invitrogen), rabbit anti-PARP1 (9542; Cell Signaling), mouse anti-PAR (4335-MC-100; Trevigen), rabbit anti-pan PAR (MABE1016; EMD Millipore), rabbit Anti-MBP (Ab9084; Abcam), mouse anti-Flag (F1804; Sigma-Aldrich), rabbit anti-macroH2A1 (Ab37264; Abcam), rabbit anti-macroH2A1.1 (12455S (DFF6N); Cell Signaling), mouse anti-macroH2A1.2 (MABE61 (14G7); EMD Millipore), rabbit normal IgG (NI01; Calbiochem), mouse normal IgG (NI03; Calbiochem), rabbit anti-ZMYND8 (A302-089; Bethyl Laboratories, Inc.).

### Immunofluorescence

For γH2AX foci and micronuclei analysis, cells were seeded onto glass coverslips and 24 h later pre-treated with DMSO or 5 μM Olaparib. After 24 h, cells were fixed with 2% (vol/vol) paraformaldehyde for 15 min, permeabilized with 0.5 % Triton X-100 in PBS for 15 min on ice and blocked with 5% BSA/PBS for 1 hour. Coverslips are then incubated with anti-γH2AX antibody (1:1000) for 1 h at RT. After 3X washes in PBS, samples were incubated with Alexa fluor 594-conjugated goat anti-mouse secondary antibody for 1 h at RT, and slides were mounted with vectashield mounting medium with DAPI. Cells were imaged using a Fluoview 3000 confocal microscope (Olympus) using a 60X objective.

### Clonogenic cell survival assay

Clonogenic cell survival was analyzed using a colony-forming assay. Briefly, 500 U2OS or HCT116 WT and KDM5A knockout cells were plated in 6-well plates 24 h before treatment. Cells were incubated with increasing concentrations of PARP inhibitor (Olaparib), and 24 h later PARPi containing media was replaced with normal DMEM media and incubated for 14 days in tissue culture incubator (37 °C, 5% CO_2_). DMSO was used as a control. Cells were pretreated with KDM5A inhibitor (CPI455) alone or in combination with Olaparib wherever indicated. Colonies were fixed and stained with crystal violet solution (0.5% crystal violet in 20% ethanol). Results were normalized to wild type untreated cells.

### Neutral comet assay

HCT116 WT or KDM5A KO cells were treated with 5 μM DMSO or Olaparib for 24 h. After treatment, DNA breaks were analyzed using CometAssay Reagent Kit (Trevigen) according to the manufacturer’s instruction. After electrophoresis, DNA was stained with SYBR-green (Invitrogen), and images were acquired with an FV3000 fluorescence microscope (Olympus). Comet tail moments (Olive) were calculated by using CometScore 2.0 software.

### X-ray irradiation

Indicated doses of Ionizing radiation (IR) were delivered by an X-ray generator (Faxitron X-ray system, RX650).

### Laser-induced live cell imaging

Laser-induced live cell imaging was performed as previously described (Kim et al., 2019a). In brief, cells were seeded onto confocal glass-bottom dishes (Wilco Wells) and were pre-sensitized by adding 10 μM BrdU for 24 h before laser-induced damage. Laser damage was induced using a 405-nm laser beam (60%) in a temperature-controlled chamber (37 °C, 5% CO_2_) using a Fluoview 3000 confocal microscope (Olympus). The relocalization intensity of GFP fused proteins at the damage sites were analyzed using FV-10 ASW3.1 software (Olympus). All images were captured using 60X oil objective lens, and recruitment intensities of GFP fused proteins were quantified by FV-10 ASW3.1 software (Olympus).

### CRISPR-mClover HR repair assay

CRISPR-mClover HR assay was performed as previously described (Kim et al., 2019b). Briefly, mClover-HR donor plasmid and Cas9-gRNA vector were transfected into U2OS WT and U2OS KDM5A KO cells. For complementation, KDM5A KO cells were transfected 24 h before mClover-HR donor plasmid and Cas9-gRNA vector transfection. After transfection, cells were incubated for 48 h, and GFP positive cells representing repair events by HR were analyzed by Accuri Flow Cytometer.

### Recombinant protein purification

U2OS cells expressing full-length GFP tagged KDM5A, KDM5B and KDM5C were harvested and lysed in high salt buffer (20 mM Tris HCl pH 7.5, 300 mM NaCl, 1 % Triton X-100 and one mM DTT). The cell lysate was incubated with GFP-trap beads for 1 h. Beads were extensively washed five times with high salt buffer, and GFP fusion protein was eluted with 200 mM Glycine pH 2.5. Eluted proteins were neutralized with 1M Tris pH 10.4. Eluted proteins were used in PAR binding assay, as described below. MBP fused KDM5A fragments were expressed in BL21 pRIL cells, purified and eluted from amylose resin with Maltose as described previously (Kumbhar et al., 2018). KDM5A 1491-1610 (F9) and KDM5A F9ΔPID were cleaved from MBP with Factor Xa and further purified using HiTrap SP HP column (GE29-0513-24) on Akta Pure FPLC (GE Healthcare).

### PAR binding assay

*In vitro* PAR binding assay was performed as previously described (Kumbhar et al., 2018). Briefly, proteins were spotted onto nitrocellulose membrane. The air-dried membrane was then blocked with 5% BSA in PBS and incubated membrane with 10 nM PAR chains for 1 h at room temperature. The membrane was washed extensively and incubated overnight with anti-PAR antibody. Immunoblot analysis was performed to detect the PAR signal. To check for PAR binding of KDM5A fragments, an equal quantity of each fragment was spotted onto the nitrocellulose membrane, and the assay was performed as explained above.

### Nascent Transcript detection at DNA damage sites

U2OS reporter cells with siRNA transfections were seeded on the coverslip in 6 well plates. To induce site-specific DSBs, U2OS reporter cells were treated with 1 μM Shield1 ligand and 1 mM 4-OHT for 4 h to mCherry-FokI expression. Transcription was induced from an integrated reporter gene by treating cells with 1 μg/ml Doxycycline for additional 3 h. Cells on a coverslip were fixed with 4% paraformaldehyde, permeabilized with 0.5% Triton X-100 in PBS, washed and mounted on glass slides with VECTASHIELD mounting medium with DAPI. Cells were visualized using Fluoview 3000 confocal microscope (Olympus).

### Bioinformatics

Algorithm Multicoil hosted at http://cb.csail.mit.edu/cb/multicoil/cgi-bin/multicoil.cgi was used for prediction of coiled-coil region in KDM5A-C (Wolf et al., 1997). Coiled-coil probability cutoff score of 0.5 and window size 28 were used to obtain dimeric or trimeric coiled coil prediction. KDM5A predicted intrinsic disorder was determined using IUPRED (Dosztanyi et al., 2005). Multiple sequence alignments were performed using ClustalOmega.

### Quantification and Statistical analyses

Graphpad Prism (version 6.0) software was used for statistical analysis. All results are presented as mean ± standard error of the mean (SEM) unless otherwise specified. P-values were calculated using unpaired t-test otherwise indicated. P-values lower than 0.05 were considered as statistically significant. Asterisks indicates P-values (*, P<0.05; **, P<0.01; ***, P<0.001, ****). N represents the number of events or independent biological replicates as described for each experiments.

### Primers

List of primers used in this study, (restriction sites in primer are underlined).

KDM5A F1: Forward: 5’-TT<u>GCGGCCGC</u>ATGGCGG GCGTGGGGCCGGGGGGCTA-3’ Reverse: 5’-TT<u>GTCGAC</u>ATCAGTGCTGAGAACCTCAGGCTC-3’
KDM5A F2: Forward: 5’-TT<u>GCGGCCGC</u>GTGCA GATGCCTAATTTAGATCTTA-3’ Reverse: 5’-TT<u>GTCGAC</u>GCTGCTTACCAGCCGCCAAAATTC-3’
KDM5A F3: Forward: 5’-TT<u>GCGGCCGC</u>CCAGTCCATATGGTTCCCACAGAA-3’ Reverse: 5’-TT<u>GTCGAC</u>ATTTACACATTGACGTCCAATGGG-3’
KDM5A F4: Forward: 5’-TT<u>GCGGCCGC</u>GCTGAAGCTGTGAACTTCTGTACT-3’ Reverse: 5’-TT<u>GTCGAC</u>CAGCTGAGCCACAGAAGCACA-3’
KDM5A F5: Forward: 5’-TT<u>GCGGCCGC</u>GCTGAGACCTGTGCTTCTGTGGCT-3’ Reverse: 5’-TT<u>GTCGAC</u>TGCTGCTGCTACCTGTGATTCCAC-3’
KDM5A F6: Forward: 5’-TT<u>GCGGCCGC</u>CGCCCTATTCCTGTGCGTCTTGAA-3’ Reverse: 5’-TT<u>GTCGAC</u>TGTCAAACACTGCAGGGCCTCTCC-3’
KDM5A F7: Forward: 5’-TT<u>GCGGCCGC</u>GTATCCCTTCAGAAGTTGCCCGTA-3’ Reverse: 5’-TT<u>GTCGAC</u>CATCATAAGTTCTTCCAGTTGTGC-3’
KDM5A F8: Forward: 5’-TT<u>GCGGCCGC</u>GAACCTCCAGTGCTGGAGTTGTCA-3’ Reverse: 5’-TT<u>GTCGAC</u>CTAACTGGTCTCTTTAAGATCCTCCAT-3’

KDM5A F8-1: Forward: 5’-TT<u>GCGGCCGC</u>GAACCTCCAGTGCTGGAGTTGTCA-3’ Reverse: 5’-TT<u>GTCGAC</u>GTCCTTTCCTTTCACTTTTAGTGG-3’
KDM5A F8-2: Forward: 5’-TT<u>GCGGCCGC</u>GAGAAACCACTAAAAGTGAAAGGA-3’ Reverse: 5’-TT<u>GTCGAC</u>CTCTTCTTCTTTTGCTAGTTTCTT-3’
KDM5A F8-3: Forward: 5’-TT<u>GCGGCCGC</u>CTGGCCAAGAAACTAGCAAAAGAA-3’ Reverse: 5’-TT<u>GTCGAC</u>GCACACAGCATTCTCATCATCAGA-3’
KDM5A F8-4: Forward: 5’-TT<u>GCGGCCGC</u>GAGGAGTCTGATGATGAGAATGCT-3’ Reverse: 5’-TT<u>GTCGAC</u>CTAACTGGTCTCTTTAAGATCCTCCAT-3’
KDM5A F8ΔPID: Forward: 5’-AAAAAGAAGGAGAAGGCTGCTGCAGCC-3’ Reverse: 5’-GTCCTTTCCTTTCACTTTTAGTGGTTTCTC-3’
KDM5A F9: Forward: 5’-TT<u>GCGGCCGC</u>GAGAAACCACTAAAAGTGAAAGGA-3’ Reverse: 5’-TT<u>GTCGGAC</u>GCACACAGCATTCTCATCATCAGA-3’

## Acknowledgments

We thank Roger Greenberg (U. of Pennslyvania) or providing the FokI DSB cell line and Hung-wen Liu (UT Austin) for providing poly(ADP-ribose) chains. We thank members of the Miller lab for helpful discussions and comments on the manuscript. Funding in Andreas Matouschek’s laboratory was provided by the National Institute of General Medical Sciences (R01GM124501) and the Welch Foundation (F-1817). Research support for work performed in Marcus Buschbeck’s laboratory was provided through grants RTI2018-094005-B-I00 and ISCIII PIE16/00011 from FEDER/Ministerio de Ciencia e Innovación - Agencia Estatal de Investigación. Funding for this project in the K.M.M. laboratory was provided by the American Cancer Society (RSG-16-042-01-DMC) and the National Institutes of Health, National Cancer Institute (R01CA1982279 and R01CA201268).

The authors declare no conflict of interest.

## Author contributions

R. Kumbhar performed the experiments unless otherwise indicated; R.Kumbhar and K.M. Miller designed and analyzed the experiments. F. Gong generated U2OS KO and stable SFB-KDM5A HEK293T cells lines; J. Perren performed KDM5A fragment cloning and laser-microirradiation of KDM5A-F8 experiments; D. Corujo and M. Buschbeck generated and characterized macroH2A-deficient and rescued HepG2 cell lines. F. Medina and A. Matouschek helped with protein purification. K.M. Miller conceived the study and supervised the project. R. Kumbhar and K.M. Miller wrote the manuscript with additional input from all other authors.

---

**Figure S1.**
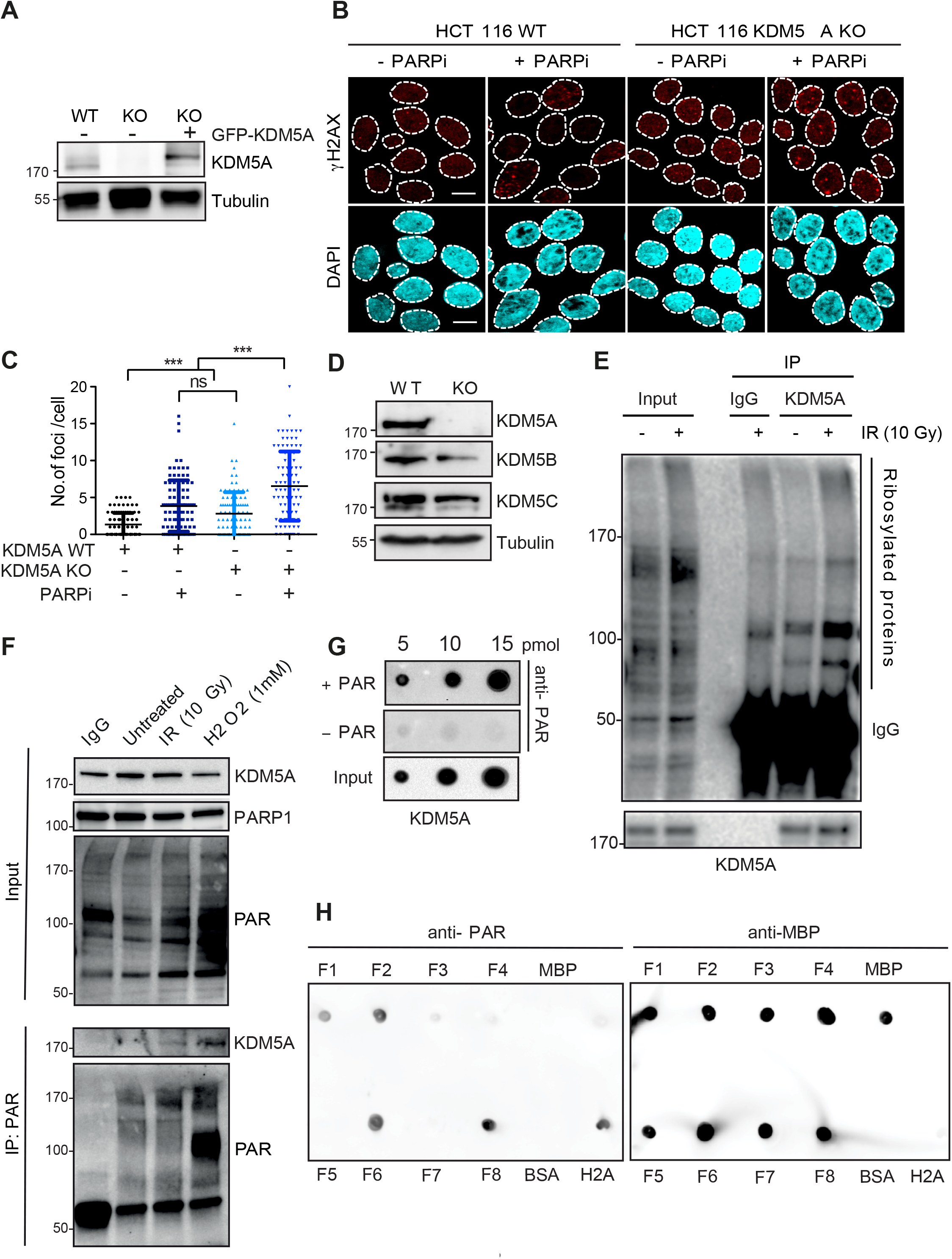
KDM5A suppresses DNA damage and interacts with poly(ADP)-ribose (Related to Figure 1 and 2). **(A)** Western blot analysis of cell lines from Fig. 1A with indicated antibodies. **(B and C)** KDM5A deficiency leads to DNA damage that is exacerbated by PARP inhibition. HCT116 WT or HCT116 KDM5A KO cells were treated ± 5 μM PARPi Olaparib for 24 h and immunostained for the DNA break marker γH2AX. γH2AX foci were quantified and plotted in B. >100 cells were quantified per condition. Error bars represents SEM. **(D)** Western blot analysis of KDM5 proteins in cells from B. **(E)** Endogenous KDM5A interacts with PAR chains. Untreated or IR treated U2OS cells were immunoprecipitated with KDM5A antibody and analyzed with indicated antibodies. **(F)** PAR IP identifies increased KDM5A interactions following DNA damage. Experiment was performed as in Fig. 2 C. **(G)** IPed KDM5A from U2OS cells binds PAR. Experiment was performed as in Fig. 2 E. **(H)** Analysis of MBP-KDM5A fragments binding to PAR. 10 pM of H2A, BSA, MBP, and MBP-KDM5A fragments F1 to F8 were spotted onto nitrocellulose and PAR binding analysis was performed as in G.

**Figure S2.**
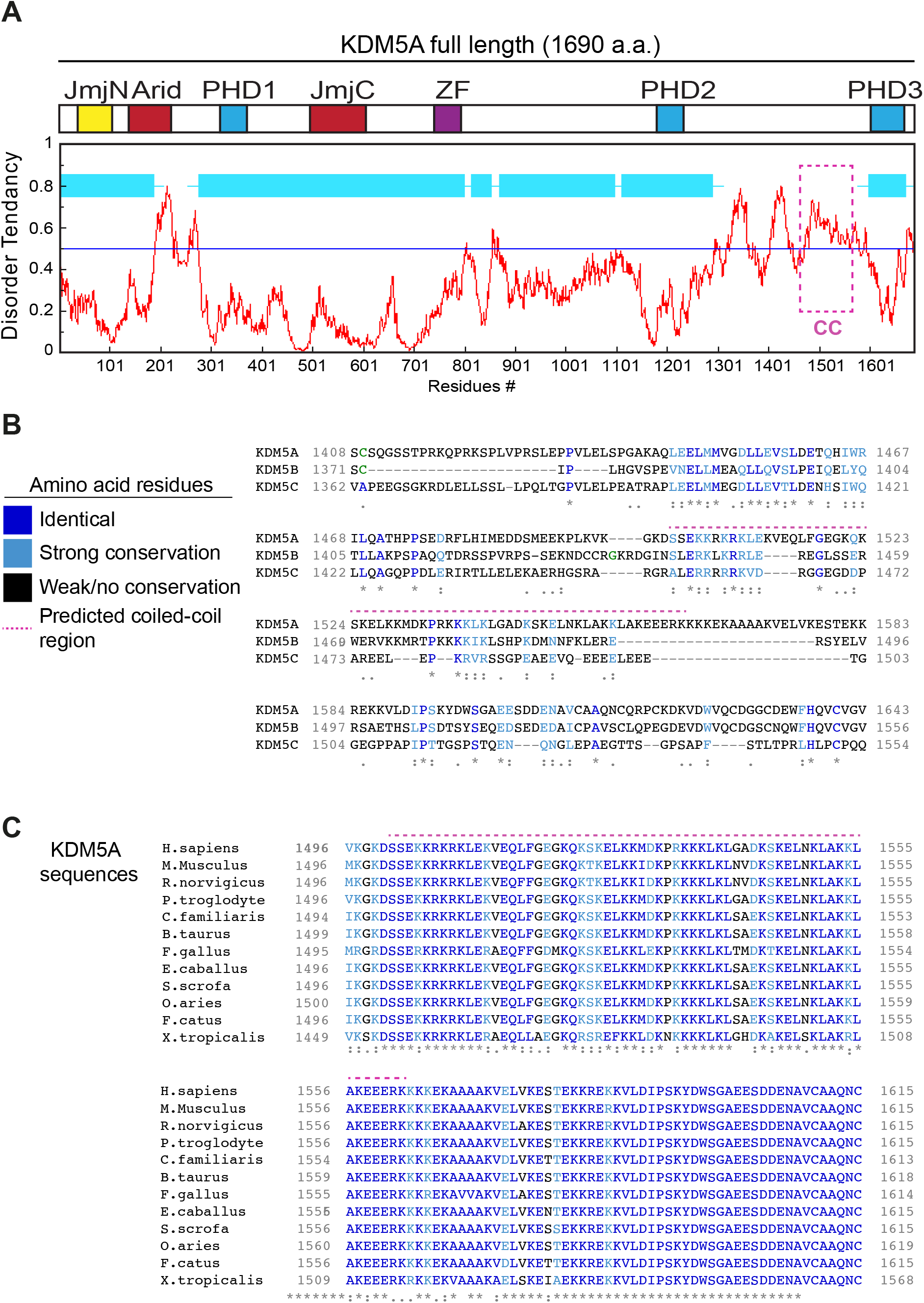
Identification of a conserved putative coiled-coil domain unique to KDM5A (Related to Fig. 3). **(A)** KDM5A PAR interacting domain (PID) is intrinsically disordered. Diagram of KDM5A aligned with a disorder tendancy plot (red graph). The disorder tendancy of KDM5A was calculated using IUPred. Globular, structured domains in KDM5A are shown in light blue boxes. Magenta box indicates PID. **(B)** Alignment of human KDM5A amino acids 1408-1623 and the corresponding regions of human KDM5B and KDM5C. Amino acids are color coordinated as indicated in the legend. PAR binding predicted coiled-coil region (CC) is indicated by the magenta dotted line. Multiple sequence alignment was carried out using Clustal Omega. **(C)** A multiple sequence alignment of *H. sapiens* KDM5A amino acids 1496-1615 with *M. musculus, R. norvegicus, P. troglodytes, C. familiaris, B. taurus, G. gallus, E. caballus, S. sacrofa, O.aries, F. catus*, and *X. tropocalis*. KDM5A exhibits high conservation in this C-terminal region. Colored amino acids are as in B. For B and C, asterisks (*) denote fully conserved residues; Colon (:) indicates conservation between strongly similar residues; period (.) indicates conservation between weakly similar residues.

**Figure S3.**
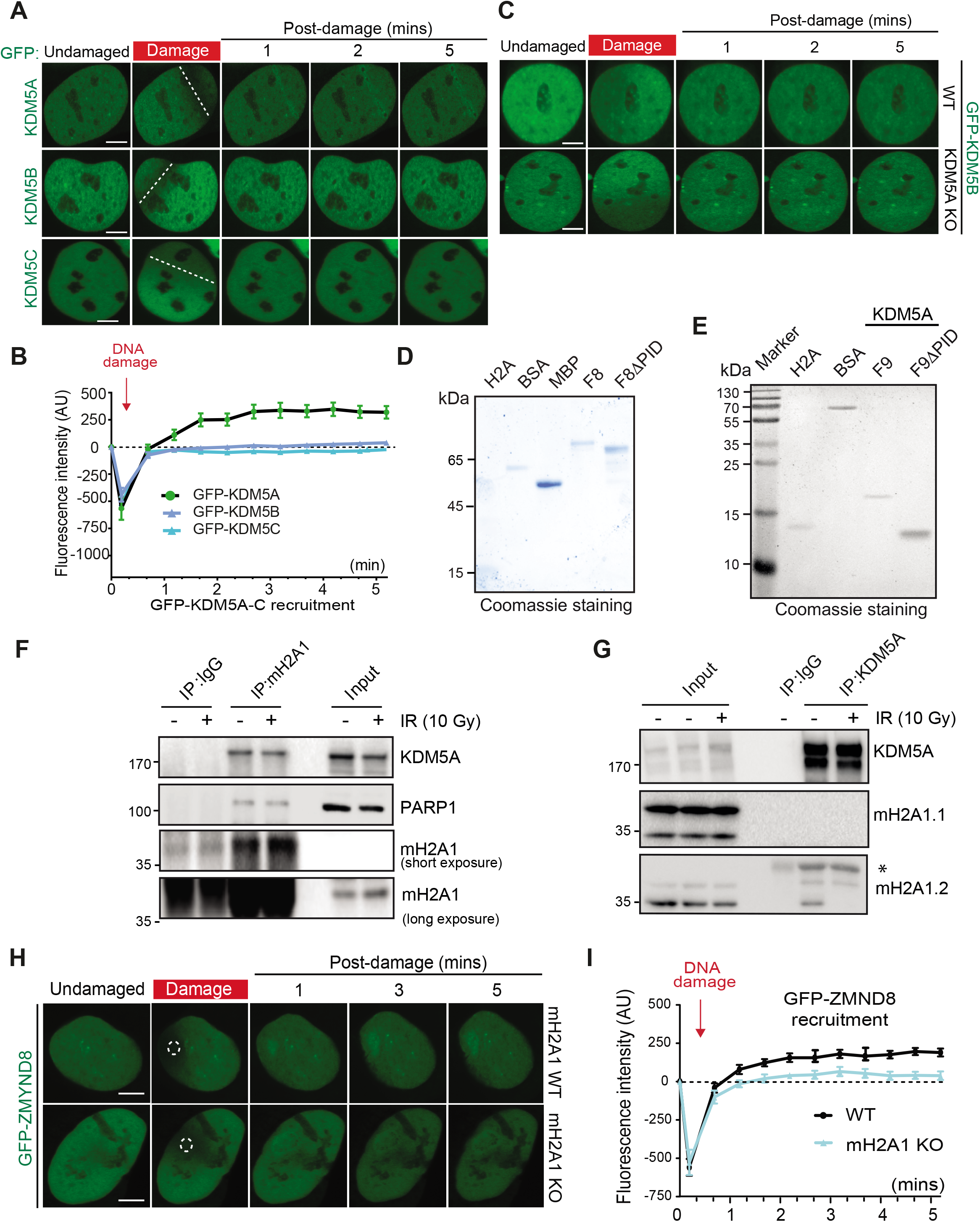
DNA damage recruitment dynamics, dependencies and interactions (Related to Fig. 3–5). **(A)** Recruitment of GFP-KDM5A, GFP-KDM5B and GFP-KDM5C to laser-induced DNA damage sites using live-cell confocal microscopy and laser-micro-irradiation. **(B)** Quantification of A. n>7. **(C)** GFP-KDM5B recruitment to laser-induced DNA damage in HCT116 WT or KDM5A KO cells. **(D and E)** Purified proteins from Fig. 3 E and G respectively were analyzed by Coomassie staining after SDS-PAGE. **(F)** Co-IP with endogenous macroH2A1 was carried out as in Fig. 2 A. **(G)** KDM5A interacts with macroH2A1.2 but not macroH2A1.1. Co-IP with endogenous KDM5A from HEK293T cells was carried out as in Fig. 2 A. **(H)** Damage recruitment of GFP-ZMYND8 is abolished in macroH2A1 KO U2OS cells. Experiments performed as in Fig. 5 A. **(I)** Quantification of H. n>10. Dotted white circles/lines indicate damaged region. Error bars represent SEM. AU, Arbitrary Units. Bars, 5 μm.

## Abbreviations

4OHT: 4-Hydroxytamoxifen
ATCC: American Type Culture Collection
AU: Arbitrary Unit
BrdU: 5-bromo-2’-deoxyuridine
CC: Coiled-coil
C-terminus: Carboxyl terminus
CRISPR: Clustered Regularly Interspaced Short Palindromic Repeats
DAPI: 4’,6-diamidino-2-phenylindole
DMSO: Dimethyl sulfoxide
DDR: DNA damage response
DSBs: DNA double-strand breaks
DOX: Doxycycline
DTT: Dithiothreitol
GFP: Green Fluorescent Protein
h: Hour
HDR: Homology directed repair
HR: Homologous recombination
IP: Immunoprecipitation
IR: Ionizing radiation
KO: Knockout
MBP: Maltose Binding Protein
NHEJ: Nonhomologous end-joining
NLS: Nuclear localization signal
N-terminal: Amino terminal
NuRD: Nucleosome remodeling and histone deacetylation
PBS: Phosphate buffered saline
PBZ: PAR-binding zinc finger
PID: PAR interacting domain
PAR: Poly(ADP-ribose)
PARPi: Poly(ADP-ribose) polymerase (PARP) inhibitor
PARGi: Poly(ADP-ribose) glycohydrolase (PARG) inhibitor
PHD: Plant homeodomain
PTM: Post-translational modification
SD: Standard deviation
SDS-PAGE: Sodium dodecyl sulphate - polyacrylamide gel electrophoresis
SEM: Standard Error of the Mean
SFB: S protein-Flag-Streptavidin binding peptide
siRNA: Small interfering RNA
shRNA: Short hairpin RNA
WB: Western blot
WT: Wild type
YFP: Yellow Fluorescent Protein

